# The dynamic emergence of musical pitch structure in human cortex

**DOI:** 10.1101/494294

**Authors:** N. Sankaran, T.A. Carlson, W.F. Thompson

**Author notes:** Current address: Department of Neurological Surgery and Center for Integrative Neuroscience, University of California, San Francisco, 675 Nelson rising Lane, San Francisco, CA 94158, USA.

## Abstract

Tonal music the world over is characterized by a hierarchical structuring of pitch, whereby certain tones appear stable and others unstable within their musical context. Despite its prevalence, the cortical mechanisms supporting such a percept remain poorly understood. The current study probed the neural processing dynamics underlying the representation of pitch in Western Tonal Music. Listeners were presented with tones comprising all twelve pitch-classes embedded within a musical context whilst having their magnetoencephalographic (MEG) activity recorded. Using multivariate pattern analysis (MVPA), decoders attempted to classify the identity of tones from their corresponding MEG activity at each peristimulus time sample, providing a dynamic measure of their cortical dissimilarity. Time-evolving dissimilarities between tones were then compared with the predictions of several acoustic and perceptual models. Following tone onset, we observed a temporal evolution in the brain’s representation. Dissimilarities between tones initially reflected their fundamental frequency separation, but beyond 200 ms reflected their status within the *tonal hierarchy* of perceived stability. Furthermore, when the dissimilarities corresponding to this latter period were transposed into different keys, cortical relations between keys correlated with the well-known *circle of fifths*. Convergent with fundamental principles of music-theory and perception, current results detail the dynamics with which the complex perceptual structure of Western tonal music emerges in human cortex within the timescale of an individual tone.

**Significance statement:** In music, pitch is organized along a hierarchy of perceived stability. Applying stimulus decoding techniques to the Magnetoencephalographic activity of subjects during music-listening, we examined the structure of this hierarchy in cortex and the dynamics with which it emerges at the timescale of an individual tone. Following its onset, we observed a temporal evolution in the brain’s representation of a tone. Activity initially reflected its pitch-value (fundamental frequency) before reflecting its status within the *tonal hierarchy* of perceived stability. ‘Transposing’ this later period of activity into different musical keys, we found that inter-key distances reflected the well-known *circle of fifths*. Our results provide a link between the complex perceptual structure of tonal music and its dynamic emergence in cortex.

## Introduction

In musical systems throughout the world, pitch is organized hierarchically (1). Depending on the prevailing *key* or *tonality* of a musical passage, certain pitch-classes occur more frequently and occupy positions of melodic, harmonic and rhythmic prominence (2). Perception mirrors this compositional hierarchy, whereby those privileged pitch-classes also have greater *stability* (3, 4). For example, within the Western key of C major, the first scale degree (C) is maximally stable and therefore heads the hierarchy. This is followed by the fifth and third scale degrees (G and E respectively), the other scale tones (D, F, G, A, B), and finally the non-scale or “out-of-key” tones (C#, D#, F#, G#, A#). We refer to this collective structure as the *standard tonal hierarchy* (STH; figure 1A).

**Figure 1.**
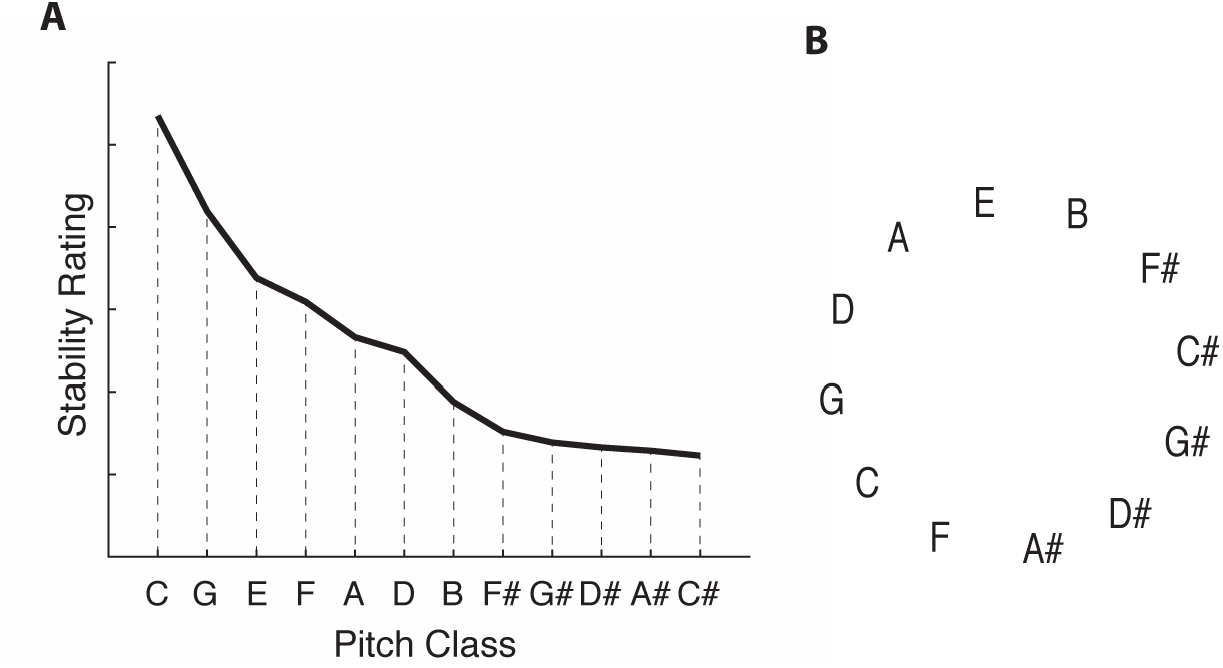
Perceptual descriptions of tonal structure. (A) The standard tonal hierarchy based on listener’s ratings of perceived stability reported in Krumhansl & Kessler (1982). (B) The “circle of fifths” conveying the relatedness between the different major musical keys.

Despite functioning as the principle organizing schema of Western Tonal Music, the neural substrates supporting the STH remain unknown. After core auditory areas extract basic frequency information from an acoustic signal, a representation of complex pitch is thought to emerge in lateral auditory regions (5-9). How does this isolated sensory representation then acquire the perceived attributes of musical pitch? The surrounding musical context must be integrated, and cortical populations reflecting a prior knowledge of Western tonal structure must be recruited. Both lesion and neuroimaging studies have identified regions implicated in the processing of both melodic (10) and harmonic (11-13) structure, while electrophysiological research has identified cortical response components sensitive to the hierarchical status of evoking tones (14-15). More recently, Sankaran et al. (2018) (16) showed that, independent from acoustics, the tonal *class* of pitches can be decoded from their multivariate patterns of Magnetoencephalographic (MEG) activity, suggesting that the perceptual structure of musical pitch may be directly recoverable from cortical activity. Despite these advances, empirical work is yet to map the neural representational space of musical pitch and explicitly test the predictions of specific perceptual and music-theoretic models. The current study therefore evaluated two major questions: Firstly, do cortical populations encode musical pitch in a manner that precipitates the organization of the STH? Secondly, what are the temporal dynamics with which afferent sensory representations of pitch interface with high-level tonal-schematic ones?

To probe these questions, we recorded the MEG activity of subjects listening to each pitch-class presented within a tonal musical context. We used Multivariate Pattern Analysis (MVPA) (17) to decode the identity of tones from their corresponding MEG activity. Within this framework, the accuracy with which classifiers can discriminate between the spatiotemporal response patterns elicited by two different tones provides an intuitive measure of their dissimilarity in cortex. As MEG responses were sufficiently time-resolved, classification was applied using a sliding time window, enabling us to track the temporal dynamics of the neural distinctions between tones. Finally, comparing the time-varying MEG dissimilarities with the predictions of relevant acoustic and perceptual models of pitch, we evaluated how the evolving cortical structure between tones relates to stimulus-driven features and the perceptual organization of the STH.

In addition to examining the brain’s representation of pitch within one key, we also measured the relationship between different major keys in cortex. This was motivated by the musical practice of *modulation*, in which a passage shifts from one key to another. In music theory, inter-key distances are described by the *circle of fifths* (figure 1B). In this arrangement, keys separated by intervals of a fifth are closest, and the pattern of relatedness folds back on itself to form a closed circle. Perceptual research has shown that these key-relations emerge when correlating the STH of different keys with one another (4), suggesting that the cognitive basis of tonality resides in the “scaffold” of individual pitch relationships rather than the general accumulation of information across a tonal passage. While prior research has investigated key-relationships using fMRI (18), the relatively poor temporal resolution prohibits an understanding of the neural mechanisms underlying the emergence of tonal structure at the timescale of an individual tone. We therefore derived a neural representation of key-distances using the measured MEG distinctions between tones. Remarkably, the extent to which two keys were related in cortex was predicted by the circle of fifths. Thus, convergent with fundamental principles in both music-theory and perception, current results provide a neuroscientific conceptualization of how complex tonal structure emerges from individual pitch-relationships within music.

## Results and Discussion

Results are derived from MEG recordings during the presentation of twelve different “probe-tones” that spanned all pitch-classes within an octave range following a C major context (see methods). Discriminant classifiers attempted to decode the MEG activity of two different tones at each time-point in the neural epoch (from −100 ms to 1000 ms relative to onset), and the resulting curve of time-varying accuracy provided a dynamic estimate of the dissimilarity in their neuronal population codes. Applying this classification procedure to every pairwise combination of the twelve different tones, we characterized the dynamic representational structure of musical pitch in cortex.

### Representational structure of musical pitch in cortex

To examine the dynamics of stimulus-specific information in cortex, we first assessed the average decoding performance when classifying all pairwise combinations of tones (figure 2A). As expected, average accuracy was at chance (50%) prior to the onset of tones (t=0) as stimulus-related information was yet to activate cortex. Following onset, neural distinctions between tones first emerged at 100 ms. Distinctions were maximal at 250 ms and remained above chance for the full extent of the neural epoch.

**Figure 2.**
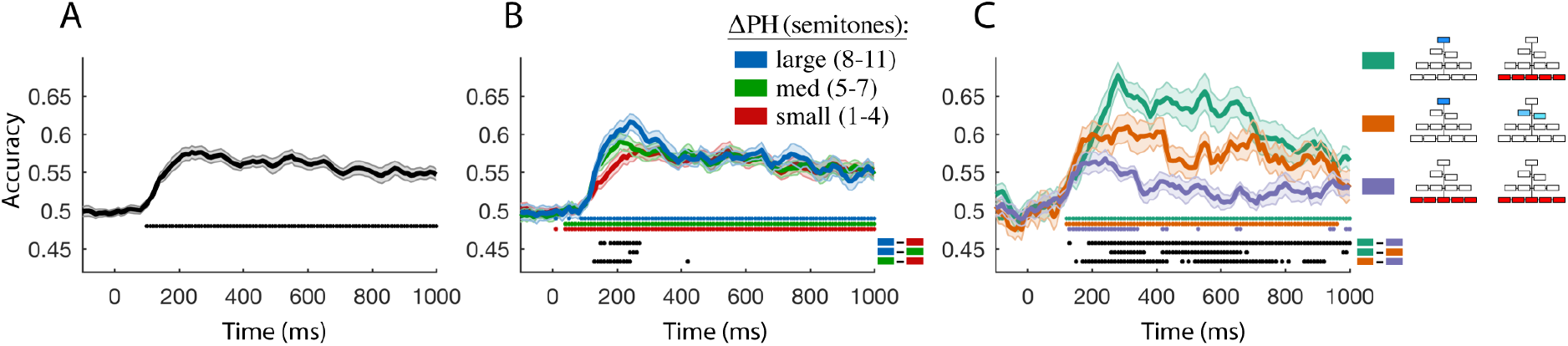
Temporal decoding of tones from evoked MEG responses. The time axis in all plots are aligned to onset of tones. (A) Average classification accuracy for decoding all pairwise combinations of the twelve tones. (B) Average classification accuracy when decoding tone-pairs grouped based on their pitch-height separation: large (8-11 semitones; blue), medium (5-7 semitones; green) and small (1-4 semitones; red). (C) Classification of tone-pairs grouped based on their difference in the hierarchy of perceived stability: large difference (green), medium difference (orange), and little-to-no difference (purple). Colored boxes in the schematic legend specify the hierarchical position of tones being decoded for each curve, with blue and red boxes indicating stable and unstable tones respectively. Results in plots B-C are averaged across all appropriate pairwise combinations of tones. Colored markers underneath curves in B-C indicate timepoints when decoding performance differs significantly from chance levels (p<0.05; Wilcoxon sign-rank tests, FDR corrected). Black markers indicate timepoints during which two decoding curves, specified by the bottom-right colored boxes, are significantly different from one another. Shaded regions indicate standard errors across all participants (N=18).

Next, we studied the dissimilarity between specific tones whose acoustic or perceptual properties generated explicit predictions regarding their representational distance. Firstly, as tones acoustically differed from one another, we reasoned that their distinctions in cortex may be commensurate with their fundamental frequency (f_0_) separation, which we term *pitch-height* (PH). Decoding performance was therefore examined for pairwise combinations of tones grouped based on whether their PH difference was small (1–4 semitones), medium (5–7 semitones), or large (8–11 semitones). A period from approximately 100 to 250 ms was found in which the above hypothesis held true (figure 2B). For example, cortical distinctions between tones that had large PH differences (blue curve) significantly exceeded those between tones that had small PH separation (red curve). Secondly, in addition to acoustic differences, tones differed in their perceived stability given the preceding musical context. We therefore hypothesized that distinctions in their cortical encoding may honor their perceptual differences, embodied by the *Standard Tonal Hierarchy* (STH) of stability. If so, MEG decoding performance would be greatest for tones located at opposite ends of the hierarchy, and poorest for tones that are hierarchically equal. In general, results confirmed this hypothesis (figure 2C). MEG responses to the most stable tone [C] were highly distinct from those of the unstable tones [F#, G#, D#, A#, C#] (green curve), but less discriminable from those of the second and third most stable tones [G and E respectively] (orange curve). Additionally, classifiers performed poorly when attempting to distinguish the neural activity of unstable tones from one another (purple curve). These results suggest that the extent to which the cortical activity elicited by two tones differ corresponds to the difference in their position within the STH. Unlike the earlier neural distinctions based on pitch-height, the correspondence between decoding accuracy and hierarchical distance only emerged approximately 200 ms after onset and persisted throughout the duration of the neural epoch.

The results of MVPA suggest that early cortical distinctions between tones reflect their absolute pitch (i.e. f_0_) differences, while later distinctions reflect the musical pitch structure of the STH. We next sought to explicitly test this hypothesis within the framework of *representational similarity analysis* (RSA) (19). The set of dissimilarities corresponding to all pairwise combinations of tones were indexed in a time-varying *representational dissimilarity matrix* (RDM; figure 3A). For a given subject and timepoint, each cell of the diagonally symmetric RDM indicates the cortical dissimilarity between the tones indexed by the cell’s row and column. We found that the RDMs of individual subjects were correlated with one another from 100 ms onwards (figure 3B), verifying that the representational structure was consistent across listeners over the same temporal extent in which average stimulus-distinctions (in figure 3A) were apparent.

**Figure 3.**
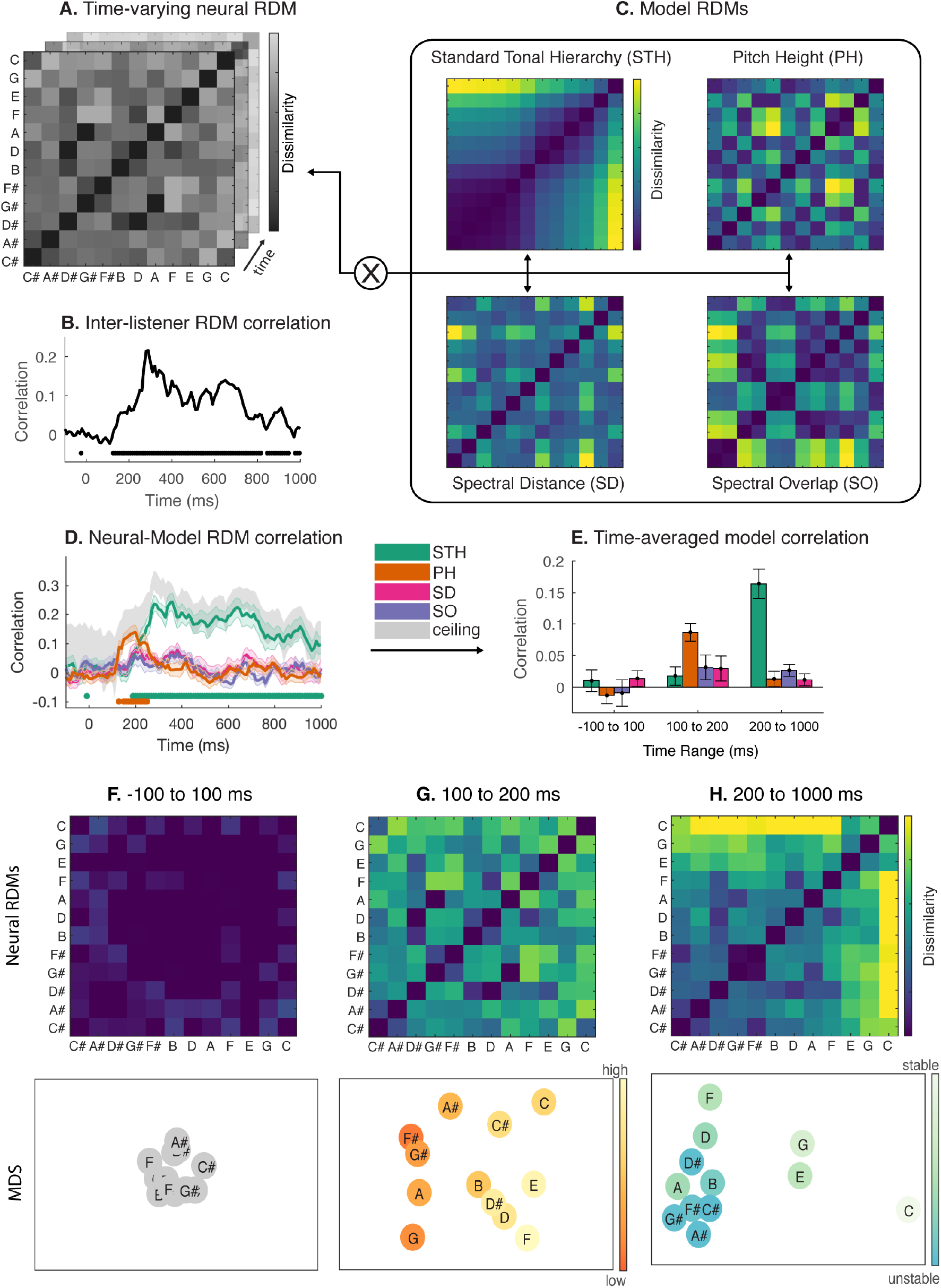
Representational similarity analysis of pitch-class. (A) Neural representational similarity matrix (RDM) indexing measured cortical dissimilarities between pairs of pitch-classes at each time point in the neural epoch. (B) The mean rank-order correlation between the RDMs of individual listeners (N=18). Significant time points are indicated underneath the curve (p<0.05; randomization test, FDR corrected). (C) Four different candidate RDMs based on models that attempt to explain neural dissimilarities. (D) Rank-order correlations between each model and neural RDMs at each time point. Shaded regions indicate standard errors across listeners. Significant time points are indicated by colored markers beneath curves (p<0.05; Wilcoxon sign-rank tests, FDR corrected). (E) For visualization purposes, neural-model RDM correlations were averaged across three different peristimulus time bins. (F-H) Average neural RDMs (top) and multidimensional scaling solutions (bottom) for the three regions in E. Colormaps indicate pitch-height (low to high) or perceived stability (unstable to stable) in G and H respectively.

Next, we evaluated the predictive capacity of several models that attempt to explain the observed structure of time-varying cortical RDMs. Each model was coded as a candidate RDM, making its predictions about the expected dissimilarities between tones explicit (figure 3C). One candidate RDM was based on the *Standard Tonal Hierarchy (STH)*, where distances between tones corresponded to their difference in perceived stability, as reported by Krumhansl & Kessler (1982) (4). Another candidate RDM coded for differences in *Pitch Height (PH)* in order to evaluate whether distinctions between tones were driven by their f_0_ separation. We additionally tested two purely sensory models; one based on the *Spectral Distance (SD)* between tone-pairs, and another based on the differences in their *Spectral Overlap (SO)* with the preceding musical context (see methods for details). Each listener’s cortical RDM at every time point was compared with the four-different candidate RDMs using a rank-order correlation measure, resulting in four curves tracking neural-model correlation across time (figure 3D). Consistent with the earlier findings from MVPA, the PH and STH models significantly explained cortical RDMs in early (100 – 250 ms) and later (190 ms onwards) regions of neural processing respectively. Crucially, both PH and STH correlations closely tracked the noise ceiling (20), indicating that these models offered optimal predictive power given the noise levels inherent in the MEG data (see methods). The temporal order of model correlations is consistent with dominant conceptions of melodic processing, which posit the extraction of complex pitch before the integration and analysis of broader tonal-harmonic structure (21). Interestingly, from 190 to 250 ms, PH and STH models were both significantly correlated with cortical RDMs, suggesting an intermediary period during which the brain holds a combined representation of both the tone’s *f_0_* and pitch-class.

To better visualize the above results, neural-model correlations were averaged into three time bins (figure 3E): the first corresponded to a period before stimulus-specific information was present in cortical activity (−100 to 100 ms); the second corresponded to the period during which cortical structure was most strongly correlated with PH differences (100 to 200 ms); and the third corresponded to the remainder of the neural epoch, during which cortical structure reflected the STH (200 to 1000 ms). Time-averaged neural RDMs corresponding to each of the three bins are displayed in the top panels of figure 3F-H. To more intuitively visualize their dissimilarity structure, we applied multidimensional scaling (MDS) to each RDM, obtaining a 2-dimensional solution in each case (Figure 3F-H; bottom panel). The MDS solution in figure 3G clearly demonstrates the organization of pitch from low to high as the space is traversed from upper-left to lower-right respectively. Similarly, the spatial organization of the MDS solution in figure 3H illustrates many key properties of the STH. Traversing the space from right to left reveals the structure of the hierarchy, with the most stable pitch-class (C) situated on the right side, closest to the next most stable classes (G and E) but distant from the cluster of unstable classes (F#, G#, D#, A#, C#) in the lower left corner. Prior behavioral research has underscored the perceptual primacy of this hierarchical arrangement (4). Our findings now provide evidence of its origins in the cortex and reveal the temporal dynamics with which it emerges from the acoustic signal via an intermediate representation of pitch-height.

### Representation of major musical keys in cortex

In tonal music, the perceptual structure that exists between individual pitch-classes is thought to generate the second-order percept of a harmonic center or “key” (22). Logically therefore, we reasoned that two keys should be related in the brain to the extent that they impose a similar neural structure amongst the constituent tones. Adapting the procedure of prior behavioral research (4) to the neural domain, we next used the MEG-based distinctions between tones to derive an empirical measure of inter-key distances, comparing the resulting structure with the *circle of fifths* (figure 1B).

In order to only capture the cortical processing of tonal-schema in our analysis, neural RDMs were first averaged across 250–1000 ms; a time during which only the STH model significantly predicted cortical RDMs. Next, we used MDS to geometrically express the dissimilarity structure between the twelve tones as points in representational space. An eleven-dimensional MDS solution was found for each subject’s neural RDM, noting that *n* objects will perfectly fit into *n-1* dimensions (23). To transpose the representational structure between tones (measured in the key of C major) into different keys, the configuration of tones was shifted in MDS-space by the appropriate number of steps entailed by a given transposition. For example, to transpose the MDS structure from C major to G major, the point representing the tone ‘C’ was shifted to that occupied by ‘G’, the point occupied by ‘C#’ was shifted to that of ‘G#’, and the process was repeated for all twelve pitch-classes. The overall dissimilarity between two keys was computed as the mean Euclidean distance across all twelve tone-translations. Application of this procedure to all pairwise combinations of the twelve major keys resulted in a cortical *inter-key* RDM in which the rows and columns correspond to different keys and cells code the corresponding distance between two keys. The average inter-key RDM across subjects is displayed in figure 4A, alongside a candidate RDM based on the circle of fifths (figure 4B). Rows and columns are ordered such that adjacent cells progress in intervals of a fifth. We found that the two structures were significantly correlated (Kendall’s Tau_A_ = 0.26; p = 0.002) and shared several essential properties. For example, keys separated by fifths were most proximate (e.g. C major and G major), while those separated by 6-semitones were most distant (e.g. C major and F# major). The generative nature of tonal music has been established by decades of perceptual research - showing that the perceived structure between keys emerges directly from the constituent structure that exists between tones (24, 25, 26, 22). Current results establish the neurophysiological basis of this generative property, deriving the same musical key-relations directly from the MEG response structure to individual tones.

**Figure 4.**
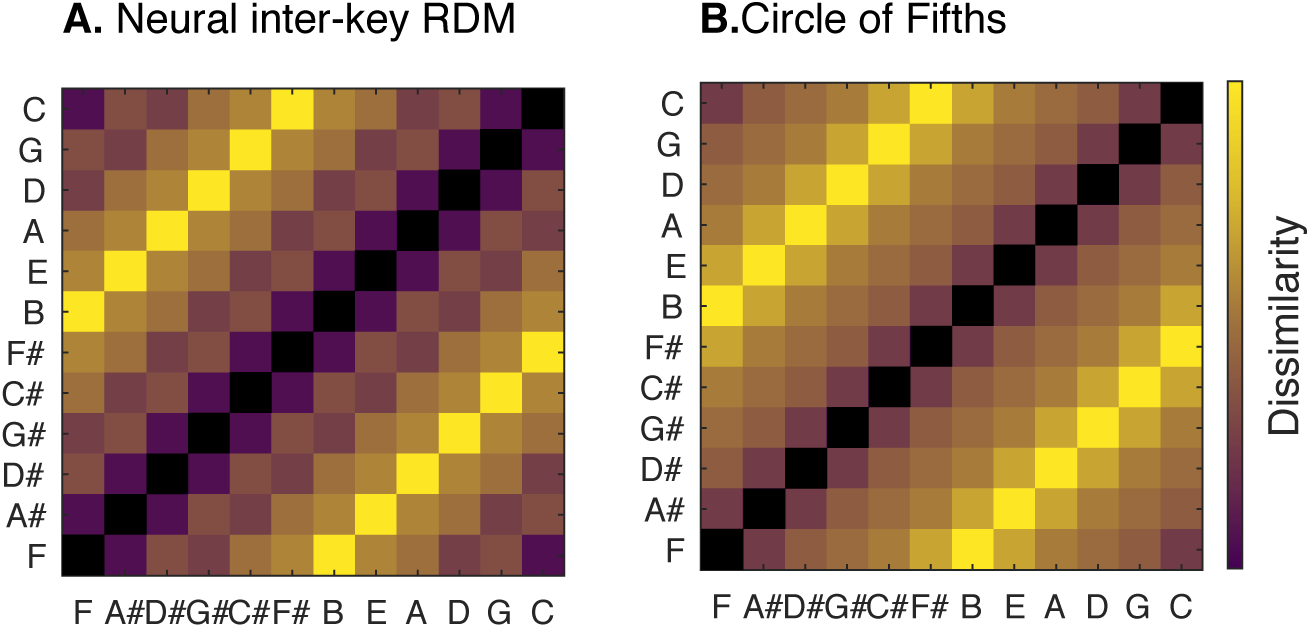
Inter-key relationships. (A) Neural RDM indicating the pairwise distances between the twelve different major keys. (B) A candidate RDM based on the “circle of fifths”.

We have dynamically characterized distinctions in the spatiotemporal patterns of cortical activity encoding the different classes of Western musical pitch. Our results suggest that, as a tone is received by the auditory system, an evolution exists in the underlying information contained in its cortical population codes. Initially, they represent a localized and intrinsic attribute of the tone. Eventually however, they contain information reflecting its integration with the surrounding context and an acquired knowledge of the pitch-structure of Western tonal music. In elucidating the representational dynamics underlying musical pitch perception, we shed light on the neural underpinnings of domain-general perceptual processes in which incoming sensory signals interact with internal structural knowledge. It remains the goal of future work to further detail the neural computations involved in integrating these two sources of information to arrive at an ultimate percept.

## Materials & Methods

### Participants

Eighteen subjects with a minimum of 5 years of formal music training (mean = 11.9 years) were recruited through the Sydney Conservatorium of Music and Macquarie University to partake in the study. All subjects reported having no known hearing loss or brain abnormalities and did not possess absolute pitch. The study was approved by the Human Research Ethics Committee at Macquarie University (REF 5201300804) and all methods were carried out in accordance with the stated guidelines. Informed consent was obtained prior to testing, after all experimental details and potential risks were explained.

### Apparatus

Data were collected with a whole-head MEG system (Model PQ1160R-N2; KIT, Kanazawa, Japan) consisting of 160 coaxial first-order gradiometers with a 50 mm baseline (27, 28). Prior to recording, each participant’s head shape was measured with a pen digitizer (Polhemus Fastrack, Colchester, VT, USA) and the positions of five marker coils on the surface of the scalp were registered. During recording, MEG data was bandpass filtered online from 0.1 – 200 Hz using first-order RC filters and digitized at 1000 Hz. Participants were in a supine position within a magnetically shielded room containing the MEG sensors. During experimental trials, they were instructed to direct their gaze at a fixation cross. Both the fixation cross and experimental instructions were projected by an InFocus IN5108 LCD back projection system (InFocus, Portland, Oregon, USA) to a screen located above the participant at a viewing distance of 113 cm. Sound stimuli were delivered via Etymonic ER-30 insert headphones at a sampling frequency of 44.1 kHz.

### Stimuli & Design

Stimuli were piano tones recorded at 44.1kHz and sampled using Max/MSP (Cycling ’74, San Francisco, CA) to construct tones that were 500ms in duration with an additional 150ms decay. Prior to testing, all tones were passed through a time varying loudness model (29) to normalize for differences in perceived loudness. For each tone, the maximum short-term-loudness (STL_max_) was computed and normalized to the mean value of all four tones. Differences in STL_max_ between all probe-tones did not exceed 3 phones.

Subjects were presented with a series of trials while having their MEG activity recorded. Each trial consisted of a tonal context followed by a single tone (hereafter referred to as the *probe-tone*). The tonal context consisted of four major chords written in four-part harmony outlining an I-IV-V-I harmonic progression in the key of C major. The context and probe-tone were separated by a silent period equivalent to one beat (650ms, 92 bpm). This temporal separation was introduced in order to prevent the sensory processing of the context from contaminating evoked responses to probe-tones whilst maintaining metric regularity. On a given trial, the subsequent probe-tone was one of twelve notes spanning the chromatic range between F#3 (185 Hz) and F4 (349 Hz). This range was chosen to minimize the average pitch-distance between the probe-tone and its preceding context. On each trial, presentation of probe-tones was randomized but constrained to avoid repeated presentation across adjacent trials. To ensure participants were attending to stimuli (30), participants judged whether the probe-tone on each trial was ‘in-key’ or ‘out-of-key’, registering their response after the occurrence of the probe-tone by pressing one of two buttons. Participants were instructed to use their left and right thumbs to register the two respective responses. The mapping of in-key/out-of-key to left/right button was interchanged every two blocks to control for the effect, if any, of motor activity on the measured neural responses. No trial-by-trial feedback was provided during the MEG recording. On average, subjects responded correctly on 78% of the trials (SD = 16.3%). All trials, including those with incorrect responses, were included in the subsequent neural analysis (31). Once the response was registered, inter-trial-intervals were randomly roved between 0.5 - 1 sec. Before testing, subjects completed a training session consisting of 20 trials with an identical behavioral task to that of the main experiment. Feedback was provided after each training trial and the experimenter ensured that subjects could perform the task (using a threshold of ≥ 75% correct) before proceeding to the MEG recording session. Each participant’s MEG data were collected in a single hour-long session. The total experiment comprised 672 trials, yielding 56 neural observations of each of the 12 probe-tones. Testing was divided into 8 blocks, each comprised of 84 trials and separated by one-minute breaks.

### Analysis

#### MEG pre-processing

Pre-processing of MEG data was performed in MATLAB. Data corresponding to each participant was first epoched from 100 ms before to 1000 ms after onset of probe-tones before being down-sampled to 100 Hz with a low-pass Chebyshev Type 1 filter. Down-sampling improved the overall SNR while still retaining a suitable level of temporal resolution to examine the time course of neural pitch-processing. Next, spatial Principal Components Analysis (PCA) was applied to the dataset of each participant using the MEG sensor channels as input features. We retained components that cumulatively explained 99% of the variance. On average, PCA reduced the dimensionality of datasets from 160 sensor channels to 28 principle components (SD = 5.4). PCA has been found to be an efficient preprocessing step for optimizing data for MEG decoding analyses (32). In a single step, PCA reduces the dimensionality of the data, and obviates the need for additional artefact rejection or de-noising procedures, as classifiers can “learn” to suppress nuisance variables isolated by PCA, e.g. eye-blinks and environmental noise.

#### Multivariate pattern classification of MEG activity

To measure the neural dissimilarity between tones, we applied Multivariate pattern analysis (MVPA) (17), whereby a binary classifier learns features of the evoked MEG activity that best distinguishes two different tones. MVPA was applied to each subject’s pre-processed dataset using MATLAB. Prior to classification, we averaged the MEG responses of 2 trials within the same pitch category in order to boost the overall SNR of classification (32). We used a naïve Bayes implementation of linear discriminate analysis (LDA) (33) to perform classification for each pairwise combination of tones. Generalization of the classifier was evaluated using k-fold cross validation with a 9:1 training to test ratio. Specifically, MEG data corresponding to the two classes being classified were randomly assigned to 10 bins of equal size, with a balanced number of observations from each class in every bin. Next, nine of the bins were pooled together and used to train the classifier, and the trials in the remaining bin were used to test the classifier. This procedure was repeated 10 times such that each bin was utilized for testing once. The reported accuracy is the average across all 10 cross-validation runs. A sliding classification time-window was used on the MEG time-series, resulting in a curve of classifier accuracy across time that tracks the dynamic emergence of stimulus-related information in the cortex. The classifier window was 50ms long and adjacent classification runs traversed the neural epoch in 10ms steps. Importantly, the neural response at each adjacent time point within the 50 ms window mapped onto a new dimension in the classification feature space. In this fashion, classifiers not only discriminated between responses based on their spatial activation patterns at each time, but also their fine-grained temporal response structure across multiple time-points. Classifier performance at each time point was evaluated in terms of balanced accuracy (32), whereby accuracy is evaluated individually for each class and then averaged. Significance at the group level (N = 18) at each time sample was evaluated using two-sided Wilcoxon sign-rank tests (p<0.05). Multiple comparisons were corrected by controlling the false discovery rate (FDR) (34, 35) with α = 0.05.

#### Representational similarity analysis of tones

Application of MVPA as described above to every pairwise combination of the twelve tones resulted in a 12×12 diagonally symmetric Representational Dissimilarity Matrix (RDM) for every subject and time sample. To check for consistency across subjects, the mean inter-subject RDM correlation (figure 3B) was calculated at each time sample by averaging the rank-order correlation (Kendall’s Tau_A_) (20) of all pairwise combinations of individual subjects’ RDMs (N=18). Significance was assessed by way of randomization testing. Briefly, the columns of subjects’ RDMs were randomly permuted before being correlated, and this procedure was repeated 1000 times, resulting in a correlation noise floor. Significance was based on the true mean correlation rising above the 95% distribution of the noise floor (FDR corrected). Next, MEG RDMs for each subject and at each time sample were compared with four candidate RDMs coded according to the predictions of several perceptual and sensory models of pitch. Candidate RDMs were as follows: [1] An RDM based on the *Standard Tonal Hierarhcy (STH)* was constructed in which each cell coded the difference in perceived stability between the two corresponding tones using the major-profile ratings reported in Krumhansl & Kessler (1982). [2] In order to test the hypothesis that MEG dissimilarities reflected the difference in each tone’s fundamental frequency (f_0_), we constructed a *Pitch-Height (PH)* RDM, in which each cell corresponded to the semitone difference in f_0_ for the two tones in question. [3] To assess whether neural dissimilarities between tones reflected their sensory differences, a *Spectral Distance (SD)* RDM was constructed in which each cell corresponded to the Euclidean distance between the 128-channel stimulus spectrograms of two tones. Spectrograms for each tone were extracted by passing the raw audio through a biologically inspired model of the auditory periphery (36). The model consisted of three main stages: (i) a cochlear filter bank comprised of 128 asymmetric filters equally distributed in log-frequency, (ii) a hair cell stage consisting of a low-pass filter and nonlinear compression function, and finally (iii) a lateral inhibitory network modelled as a first-order derivative along the tonotopic axis followed by a half-wave rectifier. (4) Lastly, although the tonal context and probe-tone were separated by 650ms (see experiment design), models of auditory short-term memory involve time-constants of up to 4 seconds (37, 38). Thus, it was possible that neural dissimilarities between tones were driven by the sensory memory of the context. To test this possibility, we constructed a *Spectral Overlap (SO)* RDM. First, the context stimulus waveform was passed through the auditory peripheral model described above in order to obtain a context spectrogram. Next, the Euclidean distance between the context and each probe-tone was calculated from their respective spectrograms. Each cell in the SO RDM was then coded as the difference in spectral distance between context and probe-tone for the two tones in question. Additional perceptual models were considered - for example the “basic space” of the *Tonal Pitch Space Theory* (25). However, the candidate RDMs arising from such models shared an identical rank-order structure to that of the STH and were therefore precluded from the analysis. Using the framework of Representational Similarity Analysis (RSA) (19), we studied the brain’s emerging representation by comparing each candidate RDM with the empirical time-varying MEG RDM (see statistical analysis below). Correlations between neural and model RDMs were assessed by computing a rank-order correlation measure (Kendall’s Tau_A_). We used the ‘noise ceiling’ as a benchmark for testing model performance (20). The noise ceiling uses inter-subject variance in RDMs to estimate the magnitude of the expected correlation between a “true” model RDM and the empirical RDM given measurement noise. To visualize the structure of RDMs, Multidimensional Scaling (MDS) was applied using Kruskal’s normalized stress 1 criterion.

#### Representational similarity analysis of musical keys

Inter-key distances in the cortex were derived by adapting the analytical approach established in Krumhansl & Kessler (1982) (4) to the neural domain. First, MEG RDMs from 250 – 1000 ms were averaged to obtain a single time-averaged neural RDM for each subject. In order to geometrically express the RDM distances between tones as points in representational space, we applied nonmetric MDS to the time-averaged RDMs of each subject. Because *n* objects will always fit into *n-1* dimensions (23), MDS solutions were obtained in eleven dimensions. Accordingly, all solutions had stress equal to zero, indicating that the MDS decomposition perfectly preserved distance information in the RDMs. To transpose the representational structure between the twelve tones into different keys, the twelve points corresponding to each pitch-class were shifted in MDS-space by the appropriate number of steps implicated by the transposition. For example, to transpose the MDS structure from C major to G major (seven semitone steps), the point representing the tone ‘C’ was shifted to that occupied by ‘G’, the point occupied by ‘C#’ was shifted to that of ‘G#’, and the process was repeated for all twelve pitch-classes. The distance between two keys was then defined as the mean Euclidean distance between the original and new positions of all twelve tones. In this fashion, distances were computed between all twelve major keys, resulting in a neural inter-key RDM for each subject, in which rows and columns correspond to different musical keys and each cell codes the corresponding distance between two keys (figure 4A). Finally, neural inter-key RDMs of each subject were rank-order correlated (using Kendall’s Tau_A_) with a candidate inter-key RDM based on the circle of fifths (figure 4B).

## Acknowledgements

This research was supported by a grant awarded to the authors from the Australian Research Council Centre of Excellence in Cognition and its Disorders, Grant CE110001021 and by a Future Fellowship awarded to TAC by the Australian Research Council, Grant FT120100816.

